# SARS-CoV-2 receptor binding mutations and antibody mediated immunity

**DOI:** 10.1101/2021.01.25.427846

**Authors:** Marios Mejdani, Kiana Haddadi, Chester Pham, Radhakrishnan Mahadevan

**Affiliations:** Department of Chemical Engineering and Applied Chemistry, University of Toronto, 200 College Street, Toronto, ON, Canada M5S 3E5; Institute of Biomedical Engineering, University of Toronto, 164 College Street, Toronto, ON, M5S 3G9

**Keywords:** COVID-19, SARS-CoV-2, Bioinformatics, Receptor Binding Domain, Mutations, Antigenic evolution

## Abstract

SARS-CoV-2 mutations can impact infectivity, viral load, and overall morbidity/mortality during infection. In this analysis, we look at the mutational landscape of the SARS-CoV-2 receptor binding domain, a structure that is antigenic and allows for viral binding to the host. We analyze 104193 GISAID sequences acquired on October 15^th^, 2020 with a majority of sequences (96%) containing point mutations. We report high frequency mutations with improved binding affinity to ACE2 including S477N, N439K, V367F, and N501Y and address the potential impact of RBD mutations on antibody binding. The high frequency S477N mutation is present in 6.7% of all SARS-CoV-2 sequences, co-occurs with D614G, and is currently present in 14 countries. To address RBD-antibody interactions we take a subset of human derived antibodies and define their interacting residues using PDBsum. Our analysis shows that adaptive immunity against SARS-CoV-2 enlists broad coverage of the RBD suggesting that antibody mediated immunity should be sufficient to resolve infection in the presence of RBD point mutations that conserve structure.

## Introduction

Coronaviruses encompass a large family of viruses that can infect both humans and animals. To date, there are seven coronaviruses that are known to infect humans, causing relatively mild to severe respiratory infections. Human coronavirus 229E, NL63, OC43, and HKU1 typically replicate within the upper respiratory tract, resulting in infections that resemble the common cold^1^. Severe Acute Respiratory Syndrome Coronavirus (SARS-CoV), Middle East Respiratory Syndrome Coronavirus (MERS-CoV), and the most recent SARS-CoV-2 can replicate within the lower respiratory tract, causing a pneumonia that can be fatal^1^.

SARS-CoV-2 is responsible for the current COVID-19 pandemic that has spread to 218 countries, infecting more than 97 million people, and causing more than 2 million deaths. This virus is most commonly characterized by fever and cough but is also associated with pulmonary embolisms, kidney injury, and gastrointestinal symptoms^2-5^. Importantly, SARS-CoV-2 has a high transmission efficiency and can lead to mortality, especially when infecting the elderly or individuals with underlying medical conditions^6,7^.

Upon entry into the respiratory tract, coronaviruses use a homotrimeric spike glycoprotein (S protein) on the surface of the virion to mediate an interaction with the host receptor angiotensin-converting enzyme 2 (ACE2) and the protease TMPRSS2^8,9^. These interactions facilitate viral envelope fusion to the cell membrane via the endosomal pathway^10,11^. The release of (+) sense RNA into the host cytoplasm allows for viral RNA translation and replication. Viral proteins and genomic RNA are subsequently assembled into virions and released from cells via vesicle transport. This infectious cycle causes host cells to undergo inflammatory pyroptotic cell death resulting in an aggressive inflammatory response and damage to the airways^1^.

The S glycoprotein is essential for the initial interaction and internalization of the SARS-CoV-2 virus by the host making it an important structure on the virion^12-15^. The S protein is comprised of the S1 and S2 domains with S1 containing the receptor binding domain (RBD) that directly interacts with the peptidase domain (PD) of the ACE2 protein (Figure 1a, b). The S2 domain contains a fusion peptide (FP), two heptad repeats, a transmembrane region, and an intracellular region (Figure 1a). RBD mediated receptor binding causes dissociation of the S1 domain and allows for the S2 domain to reach a post-fusion state enabling viral membrane fusion^10,11^.

**Fig 1:**
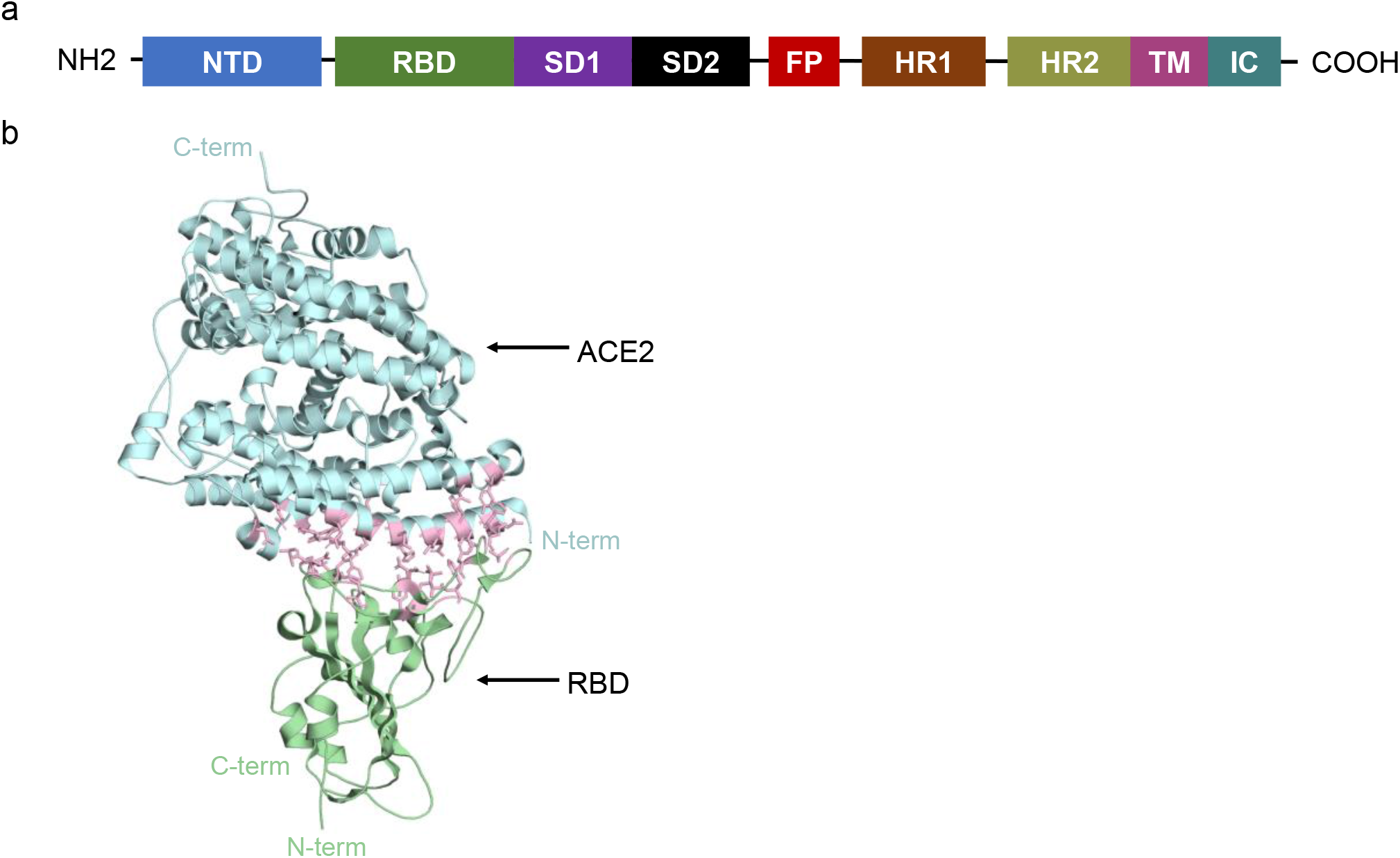
Structure of SARS-CoV-2 RBD bound to ACE2 receptor. **a**, Annotated spike monomer. NTD, N-terminal domain; RBD, Receptor binding domain; SD1 & SD2, subdomain 1 & 2; FP, fusion peptide; HR1 & 2, Heptad repeat 1 & 2; TM, transmembrane region; IC, intracellular domain. The RBD spans amino acids 330-531. **b**, SARS-CoV-2 RBD interaction with ACE2. ACE2 in cyan. RBD in green. Interacting residues are shown in pink.

Structural studies have resolved in detail the interactions between SARS-CoV-2 RBD and human ACE2, pinpointing essential contact residues (Table 1)^16-20^. Mutations in either the RBD or ACE2 can have an effect on the affinity of this interaction and may therefore impact infectivity, viral load, and overall morbidity/mortality during infection^21,22^. In this work we sought to provide insight RBD mutations using the Global Initiative on Sharing All Influenza Data (GISAID) database^23,24^. Multiple works have characterized the mutational landscape of the full SARS-CoV-2 genome or the S coding sequence using the GISAID database downloaded at various timepoints^21,25-28^. SARS-CoV-2 regions with high mutation frequency have been shown to include ORF1a, ORF1b, S, ORF3a, and N coding sequences^26^. Mutational analyses led to the discovery of a D614G mutation found on the S protein that was later shown to be responsible for higher infectivity in a pseudo typed viral infection assay, showed higher Ct values in patients, and was shown to enhance viral load in the upper respiratory tract of patients^27-33^.

**Table 1:** RBD-ACE2 Interacting Residues

| Table 1: RBD-ACE2 Interacting Residues |  |
| --- | --- |
| ACE2 Residue | SARS-CoV-2 RBD Residue |
| S19 | A475, G476 |
| Q24 | A475, G476, N487 |
| T27 | F456, Y473, A475, Y489 |
| F28 | Y489 |
| D30 | K417, L455, F456 |
| K31 | L455, F456, Y480, E484, F490, Q493 |
| H34 | Y453, L455, Q493 |
| E35 | Q493 |
| E37 | Y505 |
| D38 | Y449, G496, Q498 |
| Y41 | T500, N501, Q498 |
| Q42 | G446, Y449, Q498 |
| L45 | Q498, T500 |
| L79 | F486 |
| M82 | F486 |
| Y83 | F486, N487, Y489 |
| N330 | T500 |
| K353 | G496, N501, G502, Y505 |
| G354 | Y502, Y505 |
| D355 | T500, G502 |
| R357 | T500 |
| R393 | Y505 |

Noting the significance of these findings, it is essential that we continue to assess SARS-CoV-2 mutations as the GISAID database expands. Here we report an analysis of 104,596 GISAID RBD sequences acquired on October 15^th^, 2020. We use a combination of bioinformatics and published affinity data to determine the mutational landscape and affinities of SARS-CoV-2 RBD mutants to wild-type ACE2 proteins^21^. Finally, it has been reported that the antibody neutralization sensitivity of some RBD mutants including A475V, F490L, and V483A (among others) is reduced, suggesting that SARS-CoV-2 has mutated to evade host immunity ^27,34^. Here we analyze RBD amino acid mutations found at antibody interacting contact sites and assess the breadth of antibody mediated immunity across the RBD structure^35^. Specifically, our work focuses on mutations found at interacting residues between the RBD and the structurally characterized human antibodies BD23, COVA2-39, CV07-250, B38, CV30, CB6, P2B2F6, CV07-270, BD-368-2 and S309^35-46^.

## Results

### SARS-CoV-2 nucleotide mutations

A total of 9,275 RBD mutants were found across 104,193 sequences (8.7%) when aligned against the Wuhan reference strain (NC_045512.2). Of the 9,275 RBD mutants there were 8,871-point mutations, 179 double mutants, and 14 mutants with three to six mutations each for a total of 9064 mutant SARS-CoV-2 strains. These data are consistent with previous work showing a relatively high number of mutations at the S protein for SARS-CoV-2^26^. Analysis of mutational frequency showed that approximately 75.2% of all nucleotide mutations were G1430A, 6.8% were C1317A, 0.9% were at G1144T, and 42 additional mutants ranged between 0.1 and 0.9% frequency (Figure 2a, Supplementary Table 1). Whole genome mutational analysis of SARS-CoV-2 previously showed a high number of C -> U transitions and suggested that host-driven antiviral editing mechanisms may be driving this preference ^47,48^. In the context of RBD analysis alone, the most common mutation was a G->A transition (7.4% of mutants), followed by a C->A transversion 0.67% of the time, a G ->T transversion 0.51% of the time, and a C->U transition 0.42% of the time (Figure 2b, Supplementary Table 1). Although seemingly contrasting, these data are complementary in suggesting a high C->U transition frequency when excluding the high number of G1430A mutations in this dataset, as it represents a potentially selected mutation. Collectively, our analysis of RBD sequences show a large number of mutations occurring at the RBD with one mutant predominating the SARS-CoV-2 population.

**Fig 2:**
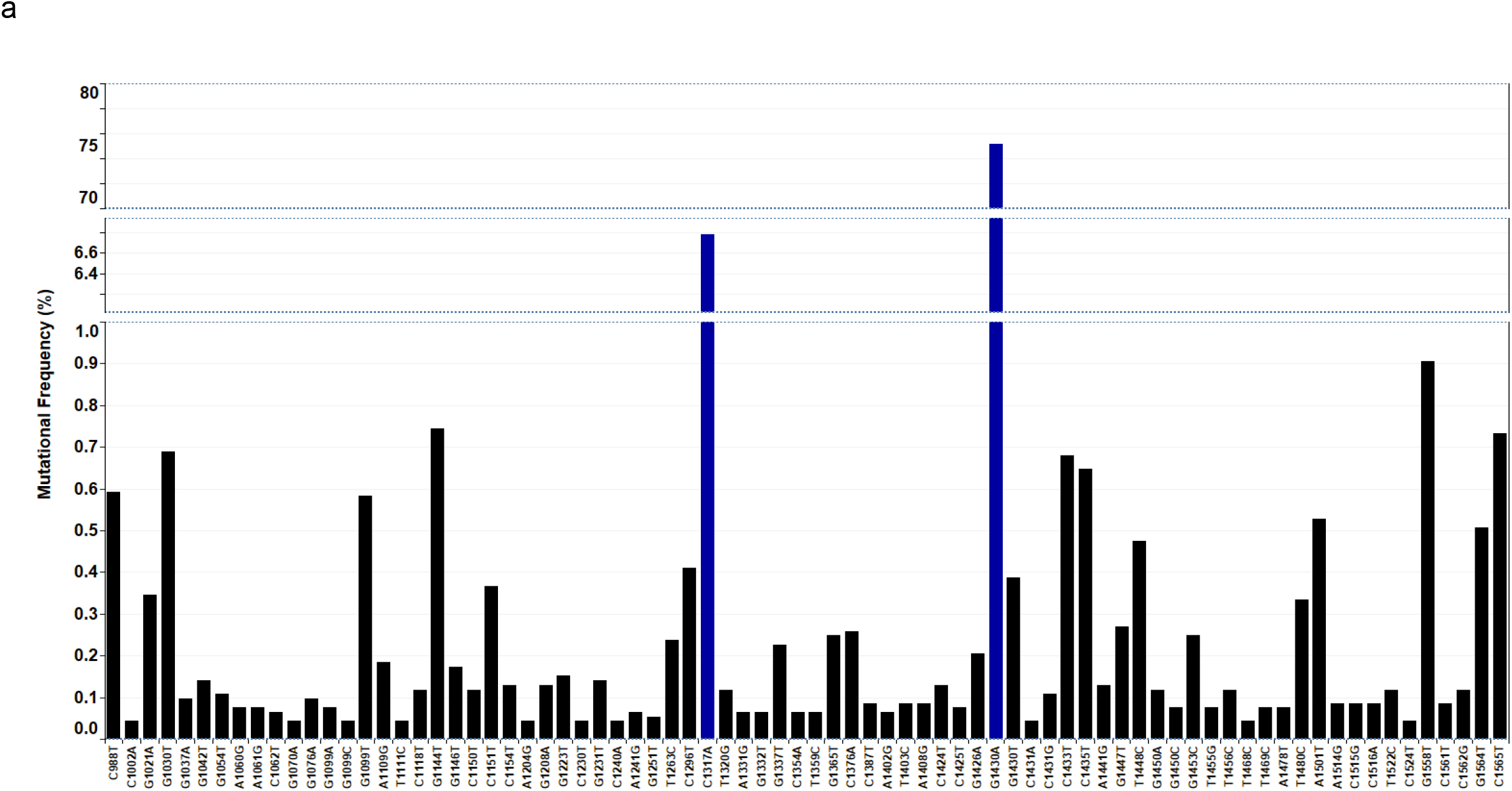
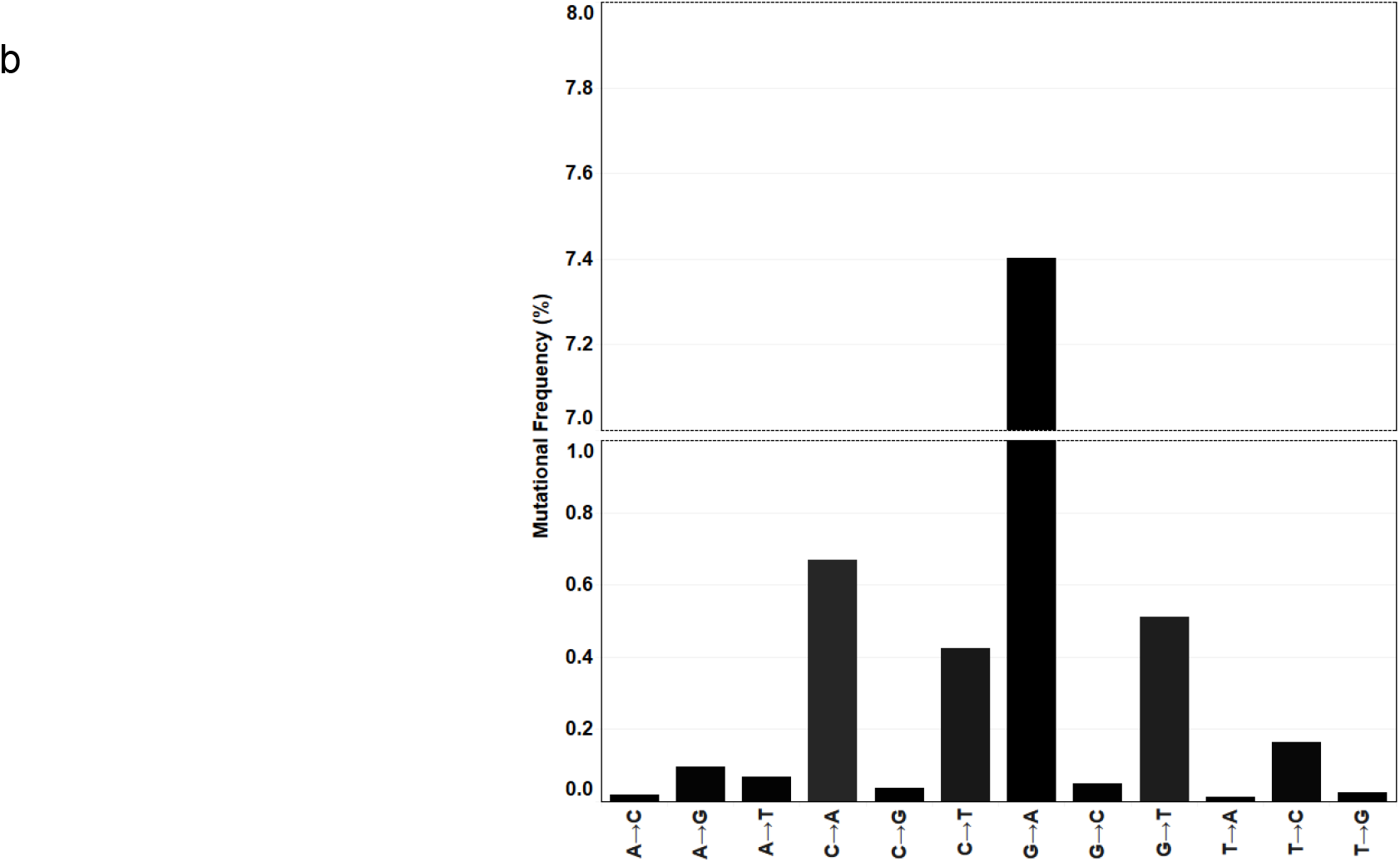
SARS-CoV-2 RBD nucleotide mutations. **a**, RBD nucleotide point mutations. Top 70 high frequency mutations across all 243 nucleotide mutations located within the RBD. Frequency reported as a function of 9064 mutant sequences. Line breaks /blue bars represent large changes in mutation frequency. **b**, Nucleotide transversion and transitions within the RBD. Frequency of transitions and transversions as a function of a total 104193 sequences.

### SARS-CoV-2 amino acid mutations

To determine the effects of these nucleotide mutations we looked at the amino acid output. Of the 9,275 nucleotide mutations approximately 1.7% of them were synonymous, whereas the remaining 98.3% modified the amino acid sequence of the RBD. The G1430A mutation led to the S477N mutant which represented 76.5% of all 9121 amino acid mutations. C1317A at 5% and G1558T at 0.9% lead to N439K and A520S mutations, respectively (Figure 3, Supplementary Table 1). Noting the high frequency of these residue modifications we wanted to assess their relevance in the context of ACE2 binding affinity. Published data on deep mutational scanning of the RBD had already reported RBD-ACE2 affinity data for all possible RBD mutants. Importantly, at the time, mutations at residues S477N and N439K represented 0.09% and 0.4% of the total GISAID sequences (31,570 total sequences)^21^. Moreover, these data showed that the S477N and the N439K mutant RBDs had a higher affinity for the ACE2 receptor than their WT counterparts, with only S477N showing improved expression in yeast cells. Here we report that these mutations now represent 6.7% (S477N) and 0.6% (N439K) of the total GISAID population (Figure 3, Supplementary Table 1). Although no significant frequency change has been observed for the N439K mutant there has been a more than 70-fold increase in the presence of the S477N mutation suggesting that selection may be playing a role in the propagation of this RBD mutant. Collectively, 13 mutations found in our GISAID analysis coded for an RBD that improved binding affinity to the ACE2 receptor, these included S477N, N439K, V367F, and N510Y amongst others (Table 2, Supplementary Figure 4). The potential negative impact of improved ACE2 binding during SARS-CoV-2 infection led us to analyze whether the high frequency S477N mutant co-occurred with the D614G spike mutation (a high frequency mutation producing more infectious SARS-CoV-2 particles). As expected, all S proteins carrying the S477N mutation also carried the D614G mutation providing evidence for this co-occurrence (Supplementary Table 2). We elaborate on this in the next section.

**Table 2.**

| RBD-ACE2 binding residue | Mutation | Count | Frequency of mutation/104193 sequences | Deep Mutational Scanning Binding Affinity<sup>21</sup> |
| --- | --- | --- | --- | --- |
| * | Y453F | 3 | 0.002879 | 0.25 |
| * | N501Y | 49 | 0.047028 | 0.24 |
| * | Y505W | 8 | 0.007678 | 0.13 |
|  | V367F | 54 | 0.051827 | 0.07 |
|  | N440K | 11 | 0.010557 | 0.07 |
|  | Y508H | 11 | 0.010557 | 0.07 |
|  | S477N | 6974 | 6.693348 | 0.06 |
| * | E484K | 11 | 0.010557 | 0.06 |
| * | Q493L | 7 | 0.006718 | 0.05 |
|  | N439K | 629 | 0.603687 | 0.04 |
|  | S359N | 9 | 0.008638 | 0.04 |
|  | N354S | 7 | 0.006718 | 0.04 |
| * | E484Q | 7 | 0.006718 | 0.03 |

**Fig 3:**
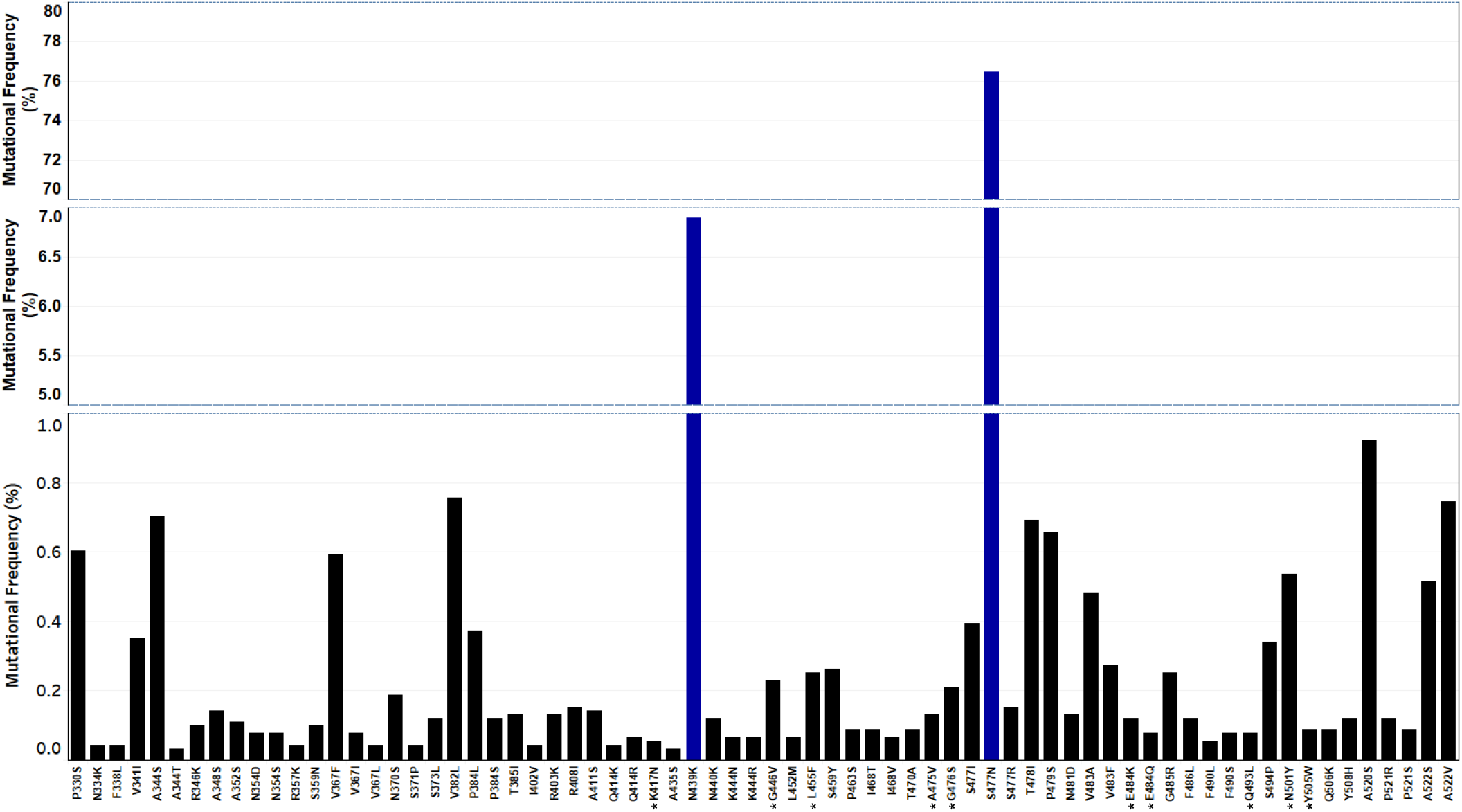
SARS-CoV-2 RBD amino acid mutations. **a**, Top 70 high frequency RBD amino acid mutations of a total 205 amino acid mutants. Line breaks/blue bars represent large changes in mutation frequency. Residues with a star represent ACE2 interacting amino acids. Frequency reported as a function of 9121 amino acid mutations.

**Fig 4:**
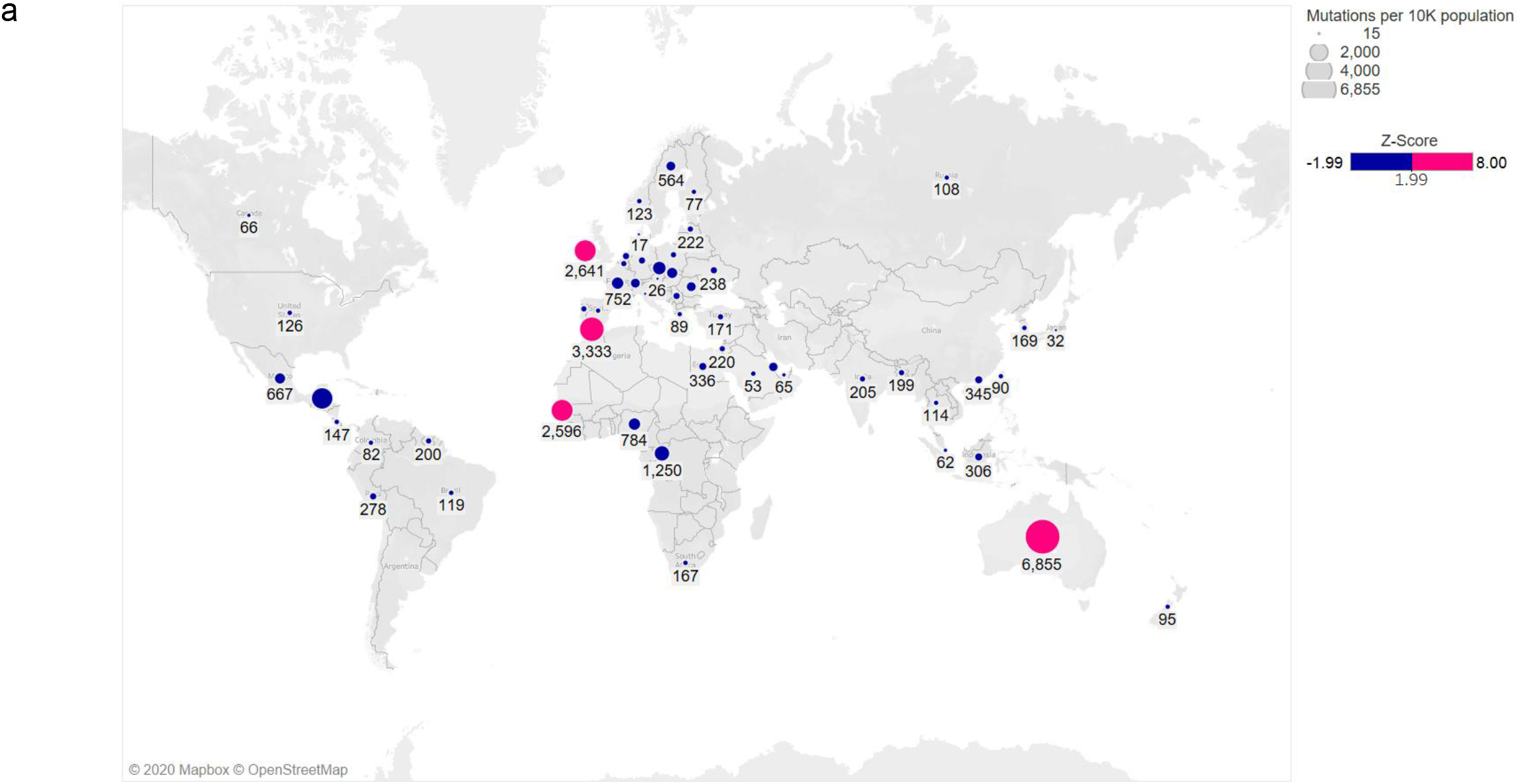
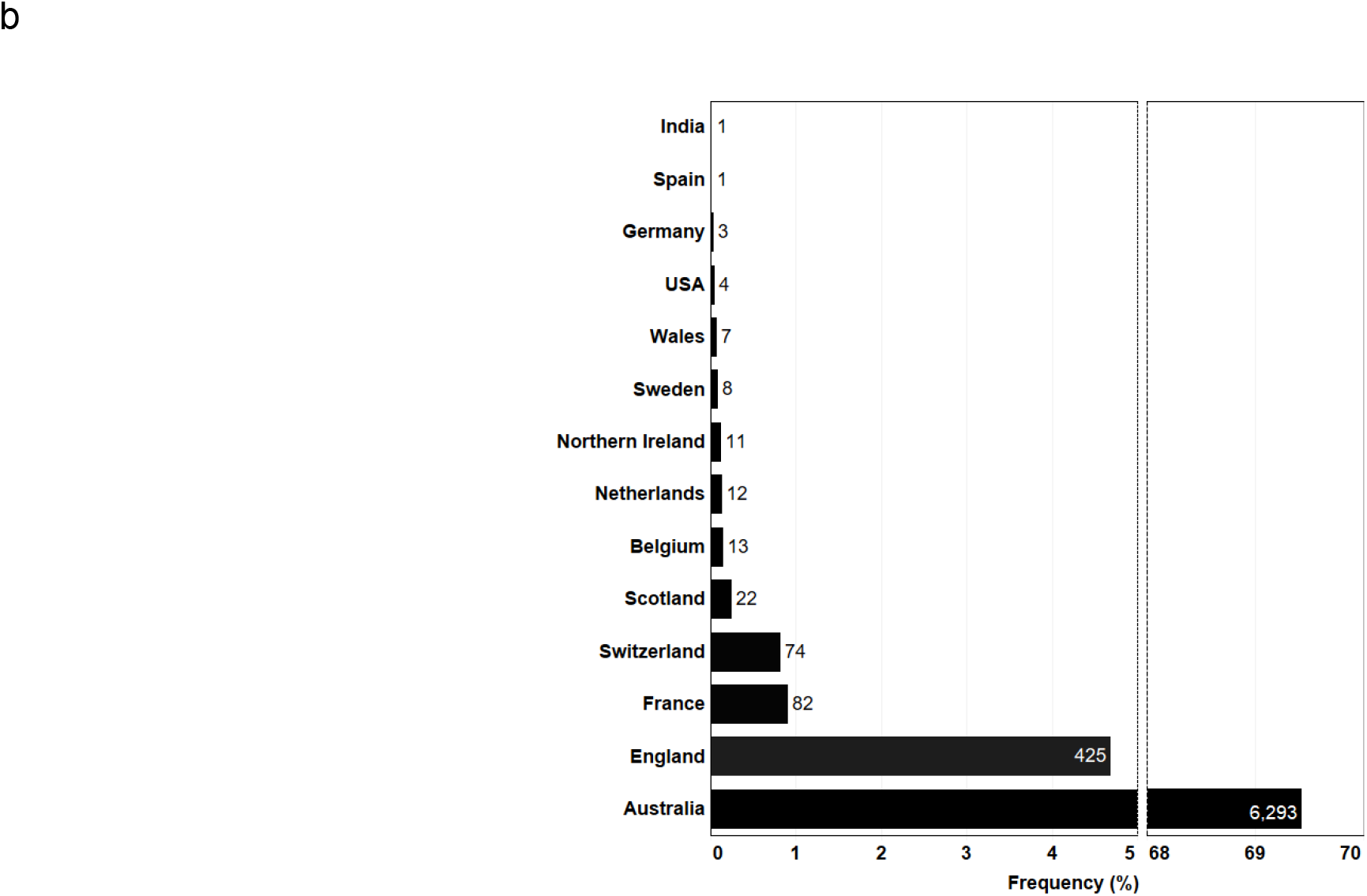
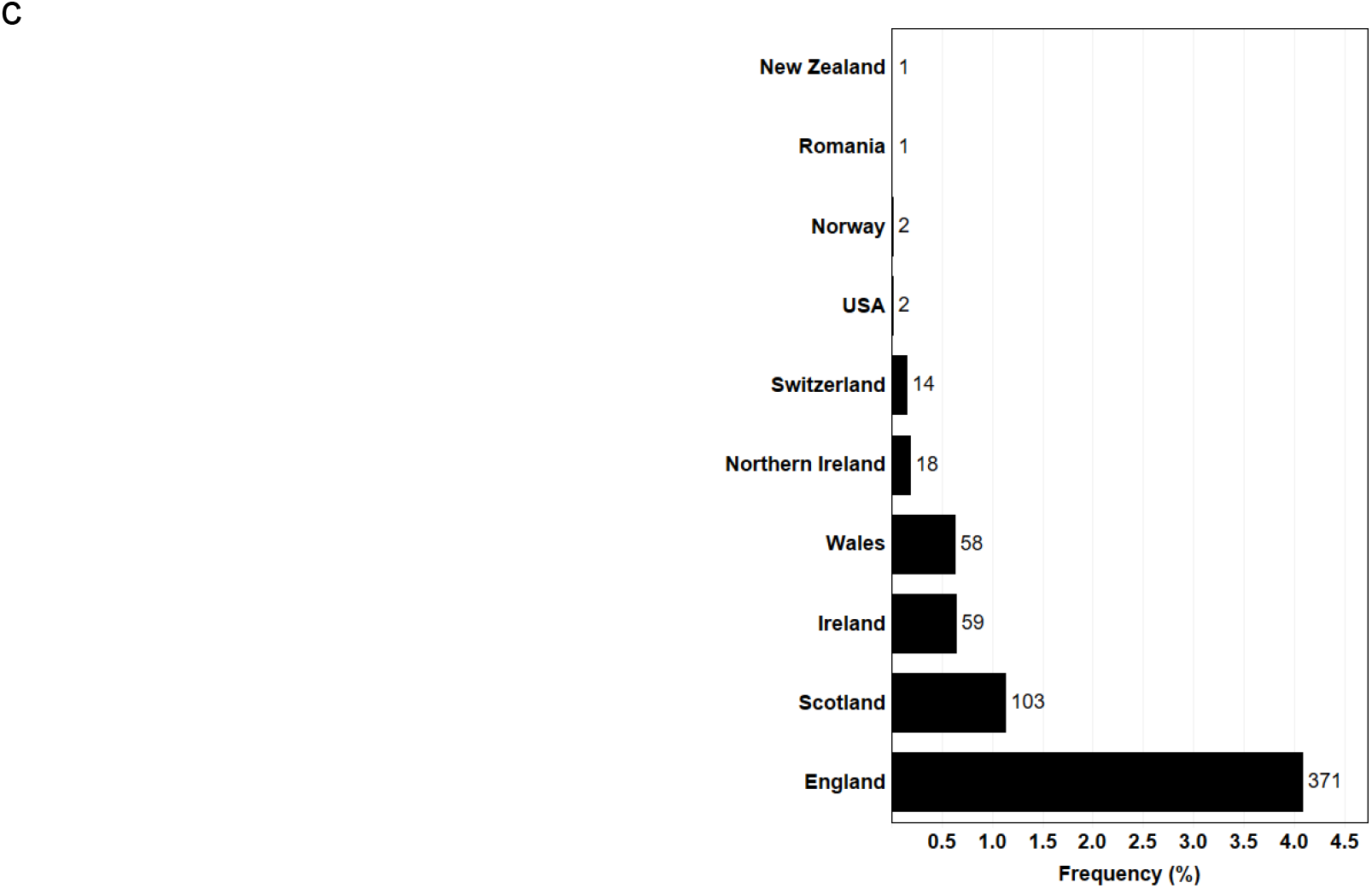
Global distribution of RBD mutations. **a**, Frequency of all RBD amino acid mutations by country. Precise Z-score values and frequencies can be found in Supplementary Figures 1 and 2, raw data located in Supplementary Table 1. **b**, Frequency of S477N mutations by country as a function of a total 6974 S477N amino acid mutants. **c**, Frequency of N439K mutations by country as a function of a total 629 N439K amino acid mutants.

### Global distribution of RBD mutations

We wanted to analyze the global distribution of individual RBD mutants as a function of country (Figure 4a, Supplementary Table 1, Supplementary Figure 1 & 2). Approximately 69% of all SARS-CoV-2 sequences in Australia contained the S477N mutation in the RBD (Figure 4b).

England, Scotland and Switzerland showed an incidence of 4.7, 0.9, and 0.8% for this mutation, respectively. The highest incidence of N439K mutants was found in England (4.1%) followed by Scotland, Ireland and Wales (Figure 4c). There was no incidence of a sequenced N439K/S477N double mutant. Overall, the S477N mutation was present in 14 countries with N439K present in 10 (Supplementary Table 2). To look at the association of D614G with S477N or N439K we reanalyzed the dataset by increasing the length of our RBD nucleotide sequence to include the D614G codon (see materials and methods for details). In both cases, all SARS-CoV-2 sequences containing either the S477N mutation or N439K mutation also contained the D614G mutation. As expected, no triple mutant was found. The S477N/D614G double mutant was found in 14 countries with a high prevalence in Australia (Supplementary Table 2). The N439K/D614G double mutant was found in 10 countries with a high prevalence in England (Supplementary Table 3).

### Mutations in antibody interacting residues found on the RBD

Structural data of human antibodies interacting with the RBD of SARS-CoV-2 was coupled with our mutational analysis. RBD mutations were found at 75% of all antibody interaction sites (Supplementary Table 5). High frequency RBD mutations included sites S477 (6.7%), V483 (0.069%), A344 (0.065%), T478 (0.065%) and N501 (0.050%) (Figure 5). S477 interacts with antibodies CV07-250, CV30, CB6, and BD-368-2, V483 interacts with COVA2-39 and P2B-2F6, A344 interacts with S309, T478 interacts with CV07-250, and N501 interacts with CV07-250, B38, and BD-368-2 (Figure 5). A number of high frequency mutants such as N439 and A520 did not show any direct interactions to antibodies in our analysis but may impact binding of other antibodies^27,34^ (Supplementary Table 5). Furthermore, no single point mutation on the RBD was found to have a direct interaction with every antibody in our analysis.

**Fig 5:**
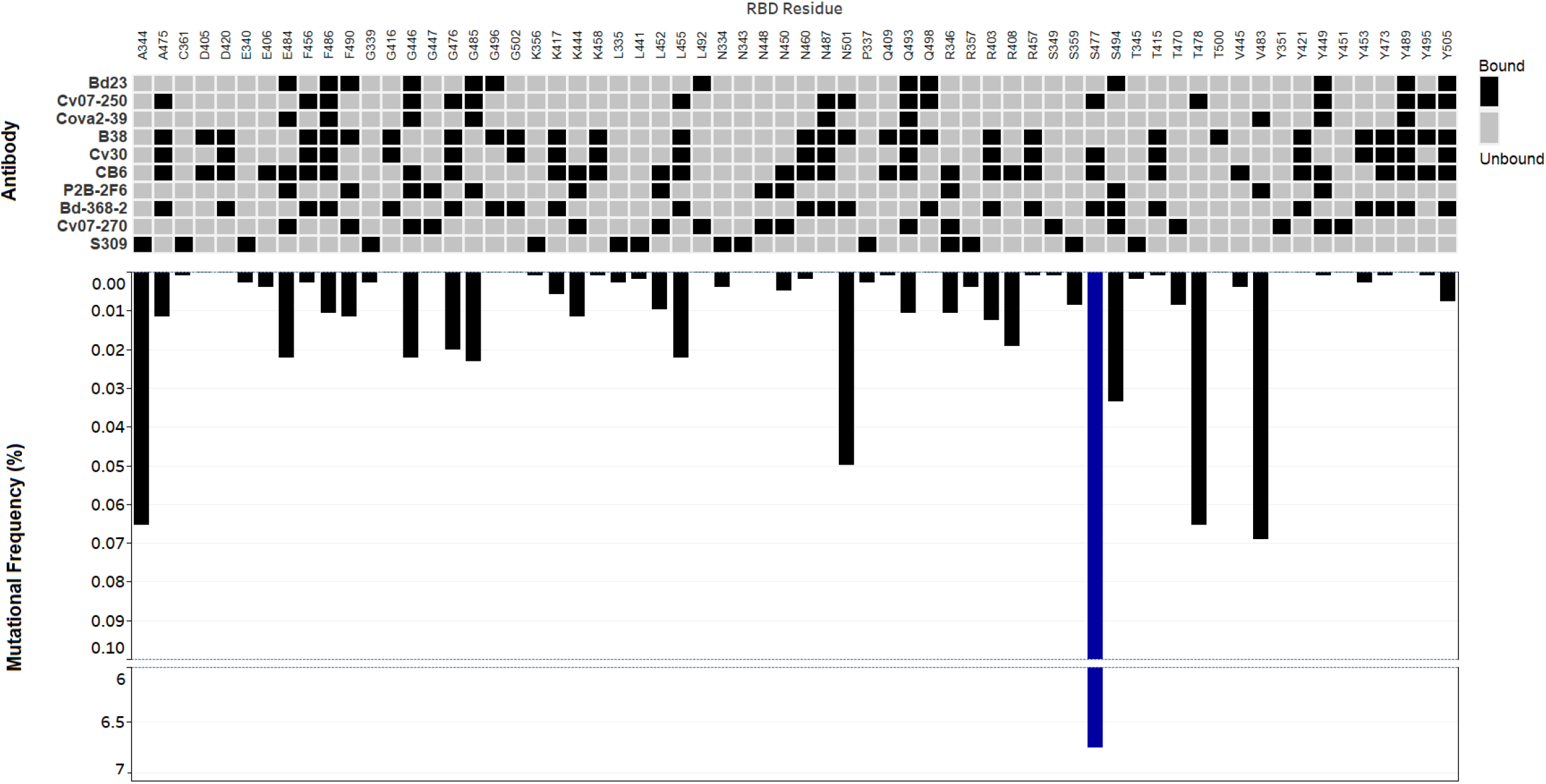
RBD mutations at antibody contact sites. Inverted bar graphs show frequency of RBD amino acid mutations as a function of 104193 sequences. Line breaks represent large changes in mutational frequency. Heat map presents antibody interactions with amino acid contact residues of the RBD.

## Discussion

Analyzing the GISAID database for SARS-CoV-2 mutations is essential as the pandemic continues. In this analysis, we provide a snapshot of all publicly available GISAID SARS-CoV-2 RBD sequences with a focus on emerging mutations that may have a negative health impact to the host. Coupling our data with previous mutational scanning work has led to multiple RBD mutants with improved affinities for their ACE2 binding target^21^ (Supplementary Table 4). Some of the highest frequency mutations with improved binding to the ACE2 receptor included S477N, N439K, V367F, and N501Y. The S477N mutation represented the highest frequency of all with N501Y displaying the highest binding affinity^21^ (Supplementary Table 4). For future analysis, it may be important to look at the potential frequency changes that occur at any RBD mutations which improve ACE2 binding affinity Table 2. The GISAID database contained no mutations at residues N487, T500, Y489, G502, and G496. This was expected as mutations at any of these key sites reduces RBD affinity for the ACE2 receptor^21^. These data strongly suggest that selection is playing a role in improving SARS-CoV-2 infectivity, with some key exceptions.

There were no SARS-CoV-2 mutations found at the Q498 residue of the RBD. This is perplexing as the Q498H, Q498Y, and Q498F all improve RBD expression and ACE2 binding, moreover, the Q498H mutation boasts the highest affinity for ACE2 of any RBD mutant^21^. Furthermore, there were a number of mutations that could significantly improve RBD-ACE2 binding but were either completely absent in our data or of extremely low frequency, including N501F, Y453F, T385R, Q493M, and Q414A amongst others^21^ (Supplementary Table 1). These contradictions may be explained by the transmission-mortality trade-off theory^49^. This theory suggests that fitness is not defined by a high mortality rate due to symptoms, but rather a high transmission rate or R_0_. It is possible that mutations at Q498 would negatively impact viral fitness by increasing mortality and reducing SARS-CoV-2 transmission. If correct, this theory suggests that the mortality rate of SARS-CoV-2 should decrease over time while transmission rate should improve to a yet unknown maximum. This may be the case with the recently increased frequency of a new spike deletion variant ΔH69/ΔV70 (unpublished work).

Finally, we expected that in some cases RBD mutations may impact the affinity of antibodies to the RBD^27,34^. To that extent we combined analysis with previously published work on RBD-antibody binding to address the potential for immune deficiencies^35^. Specifically, we analyzed ten human derived antibodies and assessed the frequency of RBD mutations at antibody contact sites. In our analysis, the pool of human derived antibodies showed that adaptive immunity against SARS-CoV-2 enlists broad coverage of the RBD. Moreover, no single RBD point mutation was found to have a direct interaction with every antibody in our analysis. This suggests that immunization with wild type and potentially any RBD point mutant that conserves structure will likely elicit the development of RBD antibodies sufficient for binding and resolving infection^27,34^. It should be noted that antibody neutralization may vary based on certain mutations as shown with the E484K mutation which in our dataset was found in 11 SARS-CoV-2 sequences^50^. The extent to which some of these mutations modify transmission efficiency and infectivity remains to be seen, but in the context of natural infection or vaccine immunization, both wild type and point mutant RBD antigens represent excellent targets for adaptive immune mediated prophylaxis.

## Materials and Methods

### Database development

A MATLAB program was developed (https://github.com/LMSE/COVID_19_Mutations) to analyze viral sequences reported on the (GISAID) database^23,24^. The GISAID database contains thousands of SARS-CoV-2 sequences isolated from patients around the world. All complete and high coverage sequences were downloaded from September 1^st^, 2019 to October 15^th^, 2020, totalling 110,761 sequences. The data was curated by removing duplicates, sequences with gaps, sequences found in animals, and sequences with a similarity score lower than 97%, yielding a 106,941-sequence dataset. The developed code generates all three reading frames for input nucleotide sequences and enables analysis of amino acid and nucleotide mutations to a reference strain. Furthermore, the code analyzes each sequence’s metadata to extract and store the collection date and the isolation location of samples reported in the database. We hope that the bioinformatic workflow and code presented here will be a valuable community tool for analyzing the spread of mutations in SARS-CoV2 sequence.

### Analyses overview

This program was used to assess the similarity of patient sequences to the reference genome of SARS-CoV-2 (NC_045512.2). Two analyses were conducted in this work. First, RBD fragment nucleotide sequences from patient samples were locally aligned to the reference genome RBD fragment, nucleotides 22544 to 23146. For the extended RBD analysis, nucleotides 22544 to 23407 were used. The RBD fragment was subsequently translated to its amino acid sequence comprising 201 amino acid residues. These RBD sequences were locally aligned to the reference SARS-CoV-2 RBD amino acid sequence. The second dataset expanded the alignment and analyzed residues 22544 to 23407 in order to include the D614 amino acid residue. The methodology for the second dataset was the same as the first. For affinity analysis, all RBD mutants with a frequency higher than 0.0067% or mutations at the RBD-ACE2 binding interface were analyzed (Supplementary Table 4).

### Sequence alignment

The program locally aligns the nucleotide sequence of patient samples and their three reading frames to a reference sequence. The algorithm executes local alignment in parallel using the MATLAB bioinformatics toolbox^51^. The alignment algorithm uses a pam250 scoring matrix to obtain the similarity scores of patient samples^52^. Similarity scores are then normalized to percentage to yield more meaningful datasets. For each sample, the reading frame with the maximum similarity score to the standard amino acid sequence is selected as a baseline for identifying mutations.

### Antibody Interacting Residues

The PDBsum server was used to obtain interacting residues on the protein-protein interface between antibodies and the RBD of SARS-CoV-2^53^. Specifically, antibodies B38 (PDB: 7BZ5), CV30 (PDB: 6XE1), CB6 (PDB: 7C01), BD23 (PDB: 7BYR), COVA2-39 (PDB: 7JMP), CV07-250 (PDB: 6XKQ), P2B-2F6 (PDB: 7BWJ), CV07-270 (PDB: 6XKP), BD-368-2 (PDB: 7CHE), and S309 (PDB: 6WPS). Detailed interactions can be found in Supplementary Table 5.

## Supporting information

Raw bioinformatics analysis

S477N-D614G co-occurence

N439K-D614G co-occurence

RBD-ACE2 affinity data

RBD-Antibody interactions

## Acknowledgments

Map data copyrighted OpenStreetMap contributors and available from https://www.openstreetmap.org. This work was supported by the Precision Medicine Initiative (PRiME) at the University of Toronto internal fellowship number PRMF2020-006. We gratefully acknowledge the authors from the originating laboratories responsible for obtaining the specimens and the submitting laboratories where genetic sequence data were generated and shared via the GISAID Initiative, on which this research is based.

---

**Supplementary Figure 1.**
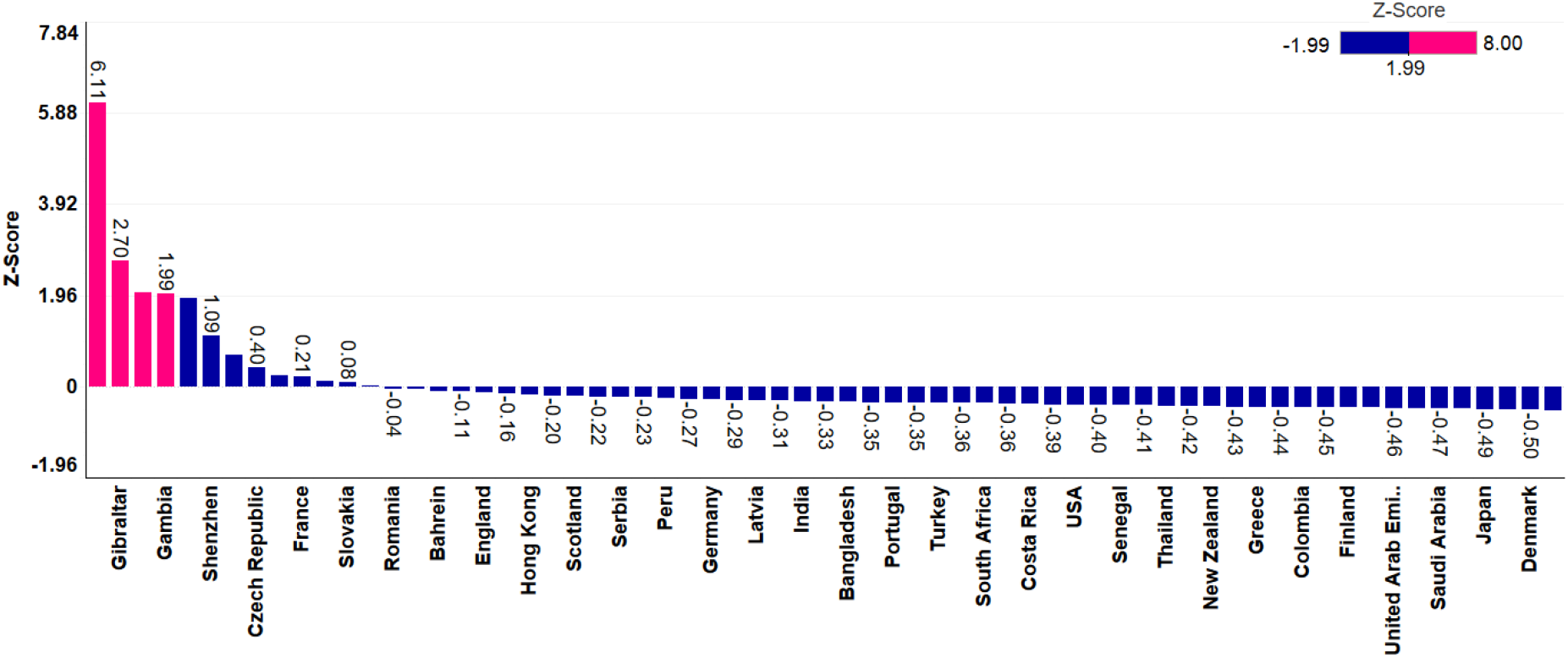
Z-scores for figure 4a. RBD mutational frequency Z-scores of total 104193 sequences per country.

**Supplementary Figure 2.**
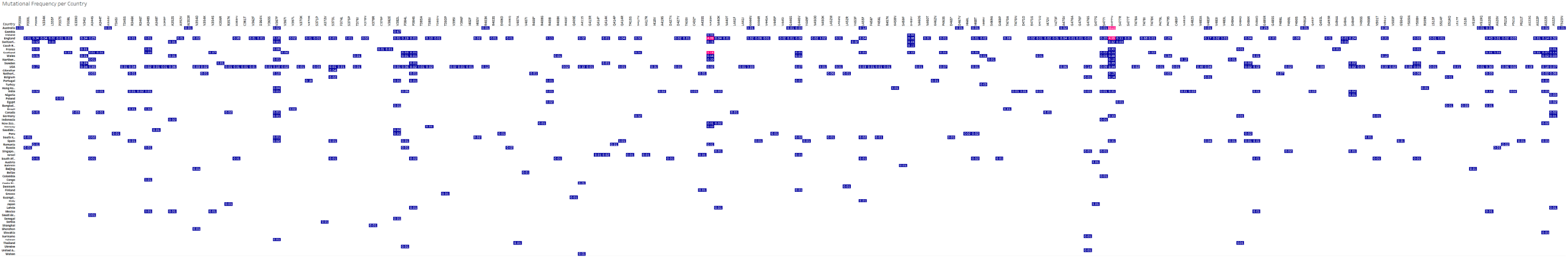
Raw mutational frequency values in percent. RBD mutational frequency of total 104193 sequences per country.

